# DCSMAP: A Hybrid Linear-Pangenome Framework for Accurate and Efficient Read Alignment

**DOI:** 10.64898/2026.10.18.759502

**Authors:** Qi Yang, Ruiwen Wan, Chun Gong, Shengkang Li, Yuxiang Li, Yong Zhang

**Affiliations:** BGI Research, Wuhan 430074, China; State Key Laboratory of Genome and Multi-omics Technologies, BGI Research, Shenzhen 518083, China

**Keywords:** short-read alignment, linear reference, pangenome reference, variant calling

## Abstract

**Background:** Linear-reference read alignment is computationally efficient but is susceptible to reference bias, whereas pangenome graph alignment better represents genetic diversity at substantially greater computational cost.

**Findings:** We developed DCSMAP, a hybrid workflow that combines linear alignment with machine-learning-based selection of reads for remapping to a personalized pangenome graph, followed by local indel realignment. An XGBoost classifier trained on linear-alignment features achieved an area under the receiver operating characteristic curve of 0.9831 on held-out test data. Across 17 runs covering seven Genome in a Bottle samples, two sequencing platforms, and two read lengths, it selected a mean of 14.80% of read pairs for graph remapping. The optimized hybrid workflows reduced variant-calling errors by 46.66% relative to BWA-MEM2 and 8.39% relative to vg personalized-pangenome alignment. Heterozygous allelic balance indicated that the hybrid workflows substantially reduced reference bias relative to BWA-MEM2 linear alignment, yielding reference bias comparable to that of the pangenome alignment. On HG003 DNBSEQ PE150 data, DCSMAP achieved a 2 to 2.5 times speedup over the personalized-pangenome workflow; both workflows used 32 threads. Efficiency gains arose from both the hybrid architecture and tool-level optimizations, which accelerated personalized-graph construction by 2.5 times, graph alignment by 1.2 times, and indel realignment by 2.5 times.

**Conclusions:** Selectively applying graph alignment to reads with potentially unreliable linear placements retains the accuracy benefits of personalized pangenomes while substantially reducing computation. DCSMAP provides an accurate and resource-efficient approach to short-read mapping on a standard CPU server.

## Introduction

Read alignment is a fundamental step in sequencing analysis and affects downstream variant discovery and genome interpretation. Most existing aligners map reads to a single linear reference, providing an efficient and well-established framework. However, a single reference cannot adequately represent genetic diversity across individuals, causing reference bias and inaccurate or missed alignments for reads carrying non-reference sequences[1–3]. Pangenome references address this limitation by representing multiple haplotypes and alternative genomic sequences, enabling more comprehensive and accurate read alignment[3,4].

Pangenome graphs provide a good representation of population-level genetic diversity, but the inclusion of numerous alternative haplotypes also increases graph complexity and the computational cost of read alignment[5]. Personalized reference approaches have therefore been developed to focus the reference representation on variation relevant to the sample being analyzed. K-mer-based methods infer sample-supported haplotypes directly from sequencing reads and construct personalized pangenome graphs, reducing irrelevant alternative paths while retaining sample-specific variation[6]. Alternatively, imputation-based approaches construct personalized diploid linear references from inferred genotypes and can further reduce reference bias while providing a more compact representation than a population pangenome[7]. Although personalization can reduce unnecessary variation in the reference, reads that are well represented by the linear reference still do not necessarily require graph-aware alignment. This observation has motivated hybrid strategies that selectively combine linear and pangenome alignment.

Several hybrid approaches have demonstrated the potential of this strategy. The Two-step Graph Mapper first estimates the most likely path through the pangenome graph and then performs linear alignment along the inferred path, reducing the search space for read placement while retaining the accuracy benefits of graph-based mapping. The authors showed that this approach achieved accuracy comparable to vg for reads containing variants while improving accuracy for reads without variants, resulting in improved overall mapping accuracy. The two-step strategy also required less total mapping time than vg, as the initial rough mapping can be performed without the graph index required for full graph alignment[8]. More recently, HiFiCCL, a reference-guided assembler for low-coverage HiFi data, optionally remaps unmapped reads by linear-reference alignment to a pangenome graph, allowing more reads to be assigned to chromosome-specific groups for downstream assembly [9]. Together, these approaches suggest that graph alignment can be selectively applied to reads that benefit from alternative haplotype information, while linear alignment remains an efficient backbone for the majority of reads.

Here, we present DCSMAP, a hybrid read alignment workflow that combines the computational efficiency of linear-reference alignment with the accuracy advantages of personalized pangenome graphs. DCSMAP first aligns reads to the linear reference with a linear-reference aligner and uses a machine-learning classifier to identify read pairs with potentially unreliable or ambiguous placements. Only these selected reads are remapped to a personalized pangenome graph. The graph and linear alignments are then integrated under one of two strategies: dcsmap applies local indel realignment only to the graph alignments before merging them with the linear alignments, whereas dcsmap-m1 first merges both sets and then applies indel realignment to the merged alignments. In addition, we optimized the principal personalized pangenome graph construction, graph alignment, and indel realignment components to improve runtime and memory efficiency. We evaluated DCSMAP across seven GIAB samples, two sequencing platforms, and two read lengths, demonstrating that it reduces variant-calling errors relative to both linear-reference and personalized-pangenome workflows while delivering a 2 to 2.5 times end-to-end speedup over the personalized-pangenome workflow.

## Results

### Overview of hybrid linear and pangenome alignment workflow

The hybrid linear and pangenome alignment workflow is presented in Figure 1. DCSMAP combines linear and pangenome alignment through machine-learning-based read partitioning. Following linear alignment, an ML classifier evaluates each read pair and partitions read pairs into those retaining their linear alignments and those selected for graph-based remapping because their linear placements are potentially unreliable or ambiguous. In parallel, KMC generates a sample-specific k-mer profile[12], which guides vg haplotypes to sample a personalized diploid graph from the population pangenome[6]. The selected reads are then remapped to this graph using vg giraffe [13] and projected onto linear reference coordinates. We evaluated two integration strategies: dcsmap realigns graph alignments and then merges the to sets, while dcsmap-m1 realigns merged linear and graph alignments. This design evaluates the effect of applying indel realignment only to graph alignments versus to the full set of graph and linear alignments, while keeping the upstream alignment procedure unchanged.

**Figure 1.**
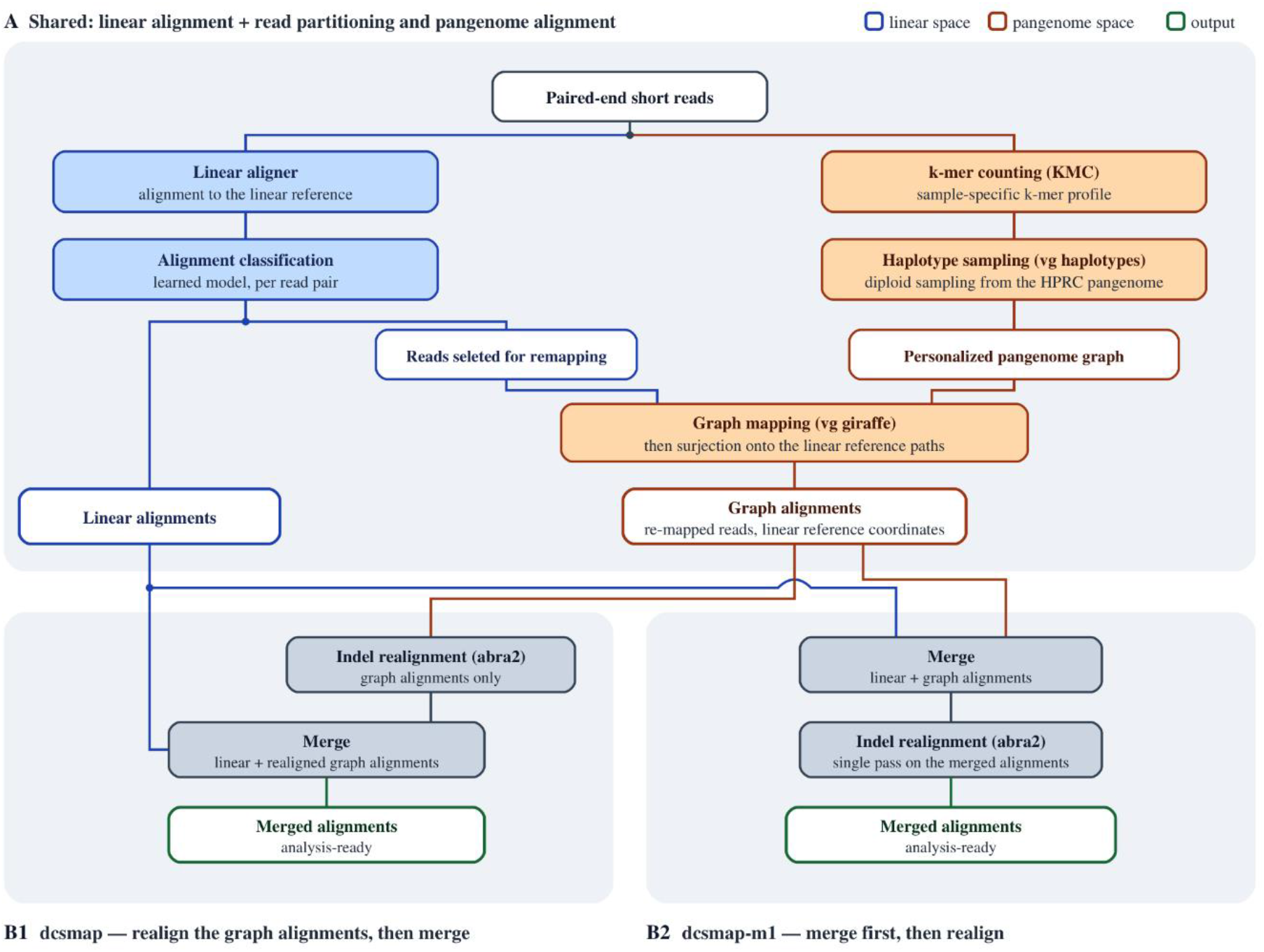
DCSMAP hybrid linear–pangenome alignment workflow. (A) Reads are first mapped to a linear reference and partitioned into reads retaining their linear alignments and reads selected for graph-based remapping because their linear placements are potentially unreliable or ambiguous. The selected reads are remapped by giraffe to a sample-specific pangenome graph constructed using KMC and vg haplotype sampling, then projected onto linear reference coordinates. (B1) dcsmap realigns graph-derived alignments with ABRA2 before merging them with retained linear alignments. (B2) dcsmap-m1 first merges both alignment sets and then performs a single ABRA2 realignment, producing analysis-ready merged alignments.

### Machine learning based read partitioning

To identify which read pairs require realignment to the pangenome graph, we trained an XGBoost classifier [14] on read-level linear alignment evidence. Labels were derived by aligning NA12891 (Illumina NovaSeq, PCR-free, 30×) with both BWA-MEM2 [15] to GRCh38 and vg giraffe [13] to a personalized pangenome subgraph and comparing the resulting placements for each read (Methods). Reads with concordant placements constituted the negative class (84.13%), whereas reads showing positional, indel, or missing-mate disagreement constituted the positive class (10.86%). Two categories, together accounting for 5.01% of reads, were excluded from model training: reads differing only in mapping quality, because a confidence difference without a positional difference does not establish which alignment is correct, and unmapped reads, which lack the alignment-derived features required by the model and are therefore unconditionally selected for graph remapping without model inference. The concordant class was downsampled to 12.5%, yielding a nearly balanced labeled dataset comprising 49.2% negative and 50.8% positive examples, which was split 80/10/10 into training, validation, and held-out test sets. Using 25 features from each read’s BAM record, the model reached an AUROC of 0.9831 and an AUPRC of 0.9853 on the validation set. At the deployed threshold of 0.50, precision was 0.9639, recall was 0.9598, and F1 was 0.9618; the held-out test set, evaluated once after all hyperparameters were fixed, agreed with validation to within 0.0002 for every metric, indicating no detectable overfitting (Supplementary Table S1).

Feature gains are strongly concentrated: IS_PROPER_PAIR (0 for not proper-pair, 1 for proper-pair) carries 69.4% of total gain and MQ a further 22.2%, so pair geometry and mapping confidence together account for 91.6% and the five highest-gain features for 97.0%. Applied in 17 end-to-end pipeline runs covering seven GIAB samples on two sequencing platforms, none of them the training sample, the classifier routed a mean of 14.80% of read pairs to graph realignment (median 14.16%, SD 2.22%, range 12.12–19.87%), from 13.23% on DNBSEQ-PE150 to 16.65% on Illumina-PE150 (Supplementary Table S2), leaving roughly 85% of the data to the linear aligner alone; that rate agrees with the 13.7% predicted from the measured recall, false-positive rate and class prevalence, demonstrating good generalizability to unseen samples, sequencing platforms, and read lengths.

### Variant-calling accuracy

Small-variant calling accuracy was evaluated for 17 Genome in a Bottle (GIAB) benchmark datasets [16] at approximately 30× coverage across two sequencing platforms (DNBSEQ and Illumina) and two read lengths (PE100 and PE150). Calls were assessed using hap.py v0.3.12 [17] against the GIAB v4.2.1 truth sets within the GIAB high-confidence regions[16,18]. Details of the evaluated workflows are provided in Supplementary Table S4. For HPRC v2.0 [19], we used the evaluate-version graph, from which HG002 and HG005 had been excluded, to avoid biasing variant-calling benchmarks toward samples whose haplotypes are already represented in the reference graph. The optimized hybrid workflows consistently achieved higher variant-calling F1 scores across all evaluated platforms (Figure 2, Supplementary Figure S1-S2, Supplementary table S5), read lengths, and samples. Averaged across dcsmap-opt and dcsmap-m1-opt, the hybrid workflows reduced the number of variant-calling errors by 46.66% relative to the BWA-MEM2 DeepVariant [15,20] workflow and by 8.39% relative to the optimized vg personalized-pangenome workflow [6,13] (Figure 3, Supplementary Figure S3, Supplementary table S8). Among the hybrid workflows, dcsmap-m1, which merges graph and linear alignments before indel realignment, achieved the highest F1 scores, at the cost of approximately 40 minutes of additional runtime for indel realignment.

**Figure 2.**
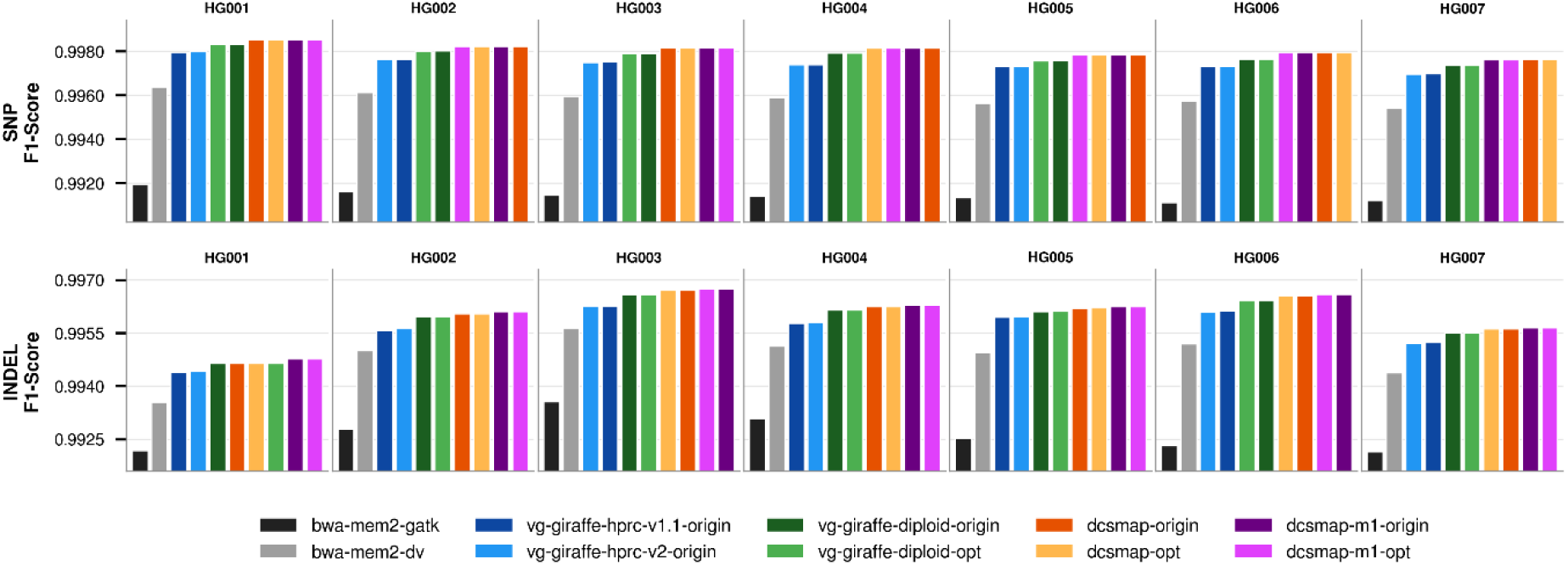
SNP and indel F1 scores on DNBSEQ PE150 data evaluated against GIAB v4.2.1. Bars show hap.py F1 scores for each workflow, separately for SNPs and INDELs across seven GIAB samples (HG001–HG007) within the GIAB v4.2.1 high-confidence regions. Within each panel, workflows are ordered by ascending F1 score. The hybrid workflows consistently achieved the highest F1 scores, with dcsmap-m1 generally providing the best accuracy. The nearly identical scores of matched original and optimized workflows indicate that the computational optimizations did not affect variant-calling accuracy.

**Figure 3.**
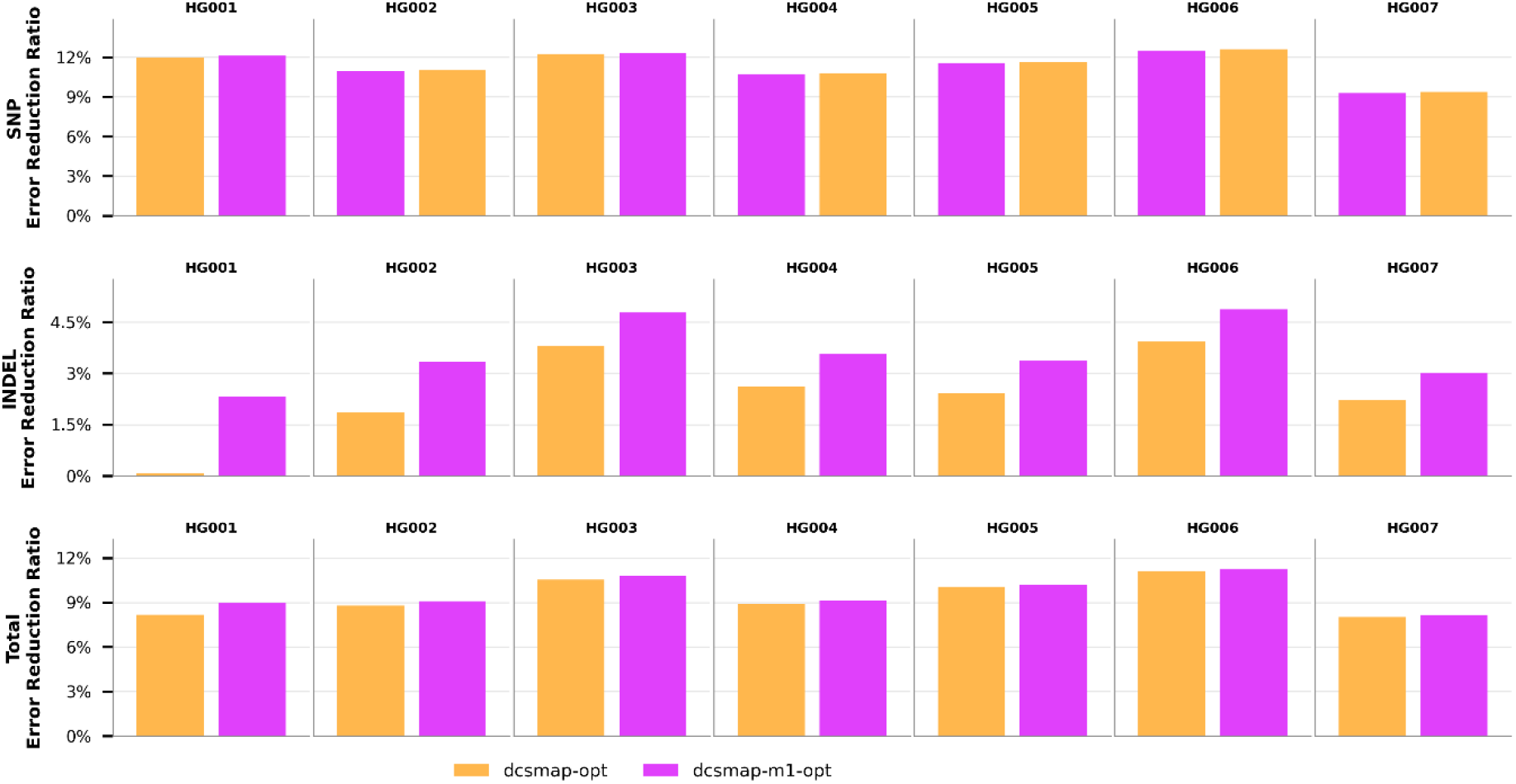
Reduction in SNP and INDEL calling errors relative to the personalized-pangenome workflow(vg-giraffe-diploid-opt) on DNBSEQ PE150 data. Total errors were calculated as the sum of false-positive and false-negative calls reported by hap.py within the GIAB v4.2.1 high-confidence regions.

To determine whether the computational optimizations made to the vg pangenome toolkit [3] and ABRA2 indel-realignment [21] software affected variant-calling accuracy, we compared three pairs of original and optimized workflows: dcsmap-origin versus dcsmap-opt, dcsmap-m1-origin versus dcsmap-m1-opt, and vg-giraffe-diploid-origin versus vg-giraffe-diploid-opt. Across 102 paired comparisons (17 datasets x 3 workflows x 2 SNP/INDEL variant types), the differences in F1 score between the original and optimized implementations were centered near zero and were not statistically significant (paired t-test, P = 0.4483, Supplementary table S10). The optimized workflows had higher F1 scores in 40 comparisons, the original workflows performed better in 42, and 20 comparisons were tied (Supplementary table S11). These results indicate that the computational optimizations did not alter variant-calling accuracy.

Across the 17 datasets, dcsmap-opt produced an average of 8.09% (Supplementary table S8) fewer variant-calling errors than the optimized personalized-pangenome workflow. To investigate the basis of this improvement, we compared dcsmap-opt (hybrid alignment) with vg-giraffe-diploid-opt (personalized-pangenome alignment) using DNBSEQ PE150 data for HG003; false negatives accounted for 87% of the error difference, and we identified 1,793 false-negative sites unique to vg-giraffe-diploid-opt. At these sites, vg-giraffe-diploid-opt had a lower median read depth than dcsmap-opt (16 versus 32), and 198 sites (11.0%) had no read. Its median per-site fraction of reads with MAPQ ≥ 60 was also substantially lower (0 versus 1.00) (Supplementary Figure S4), while most dcsmap reads at these loci originated from the linear aligner BWA-MEM2 (median, 97.8%; mean, 79.8%). Thus, although vg-giraffe-diploid-opt aligns reads to a sample-specific personalized pangenome graph, redundant or homologous sequence paths remain in the graph and may create competing placements, leading to lower MAPQ or displacement of reads away from the target locus and consequently reducing variant support and DeepVariant sensitivity.

For the HG002 sample, we repeated the benchmarking experiments described above and evaluated the results against the HG002 v5.0q benchmark set, which was constructed using a haplotype-resolved assembly-based approach and provides more comprehensive variant calls in complex and previously challenging genomic regions than GIAB v4.2.1 [22]. Consistent with the results observed on GIAB v4.2.1, the hybrid workflows achieved the highest F1 scores on both platforms (SNP, 0.9945–0.9948; indel, 0.9672–0.9758), exceeding the vg personalized pangenome workflow (SNP, 0.9944–0.9947; indel, 0.9667–0.9755) and the BWA-MEM2 DeepVariant workflow (SNP, 0.9912–0.9916; indel, 0.9610–0.9734) (Supplementary Figure S5-S6, Supplementary Table S6). Within the HG002 CMRG regions [23], the hybrid workflows achieved F1 scores slightly below the vg personalized-pangenome workflow (SNP, 0.9865–0.9883 versus 0.9874–0.9889; indel, 0.9680–0.9706 versus 0.9711) but substantially above BWA-MEM2–DeepVariant (SNP, 0.9757–0.9765; indel, 0.9589–0.9600) (Supplementary Table S7).

### Allelic balance at HETs

To obtain a direct, read-level measure of reference bias, we assessed allelic balance at heterozygous (HET) sites using biastools [24]. HET sites were taken from the GIAB v4.2.1 benchmark genotypes and analyzed within the corresponding high-confidence regions[18], and allelic balance, defined as the fraction of ALT-supporting reads among all reads covering a site, was summarized for variants stratified by allele length, with SNPs shown at the center, deletions to the left, and insertions to the right (Methods). On DNBSEQ PE150 data for HG003 (Figure 4), the linear-reference workflow (BWA-MEM2) exhibited the strongest reference bias, departing most markedly from the expected 0.5 balance for the longer insertions and deletions at the two extremes of the allele-length spectrum. The hybrid alignment workflow yielded allelic ratios closer to the expected 0.5:0.5 than BWA-MEM2 across variant lengths, indicating reduced reference bias, and allelic balance for the dcsmap-m1 workflow was closely comparable to that of the pangenome graph-based alignment methods. Similar trends were observed on data from the other two sequencing configurations, namely Illumina PE150 data for HG003 (Supplementary Figure S7) and DNBSEQ PE100 data for HG001 (Supplementary Figure S8), indicating that the reduction in reference bias achieved by the hybrid workflow is independent of sequencing platform and read length.

**Figure 4.**
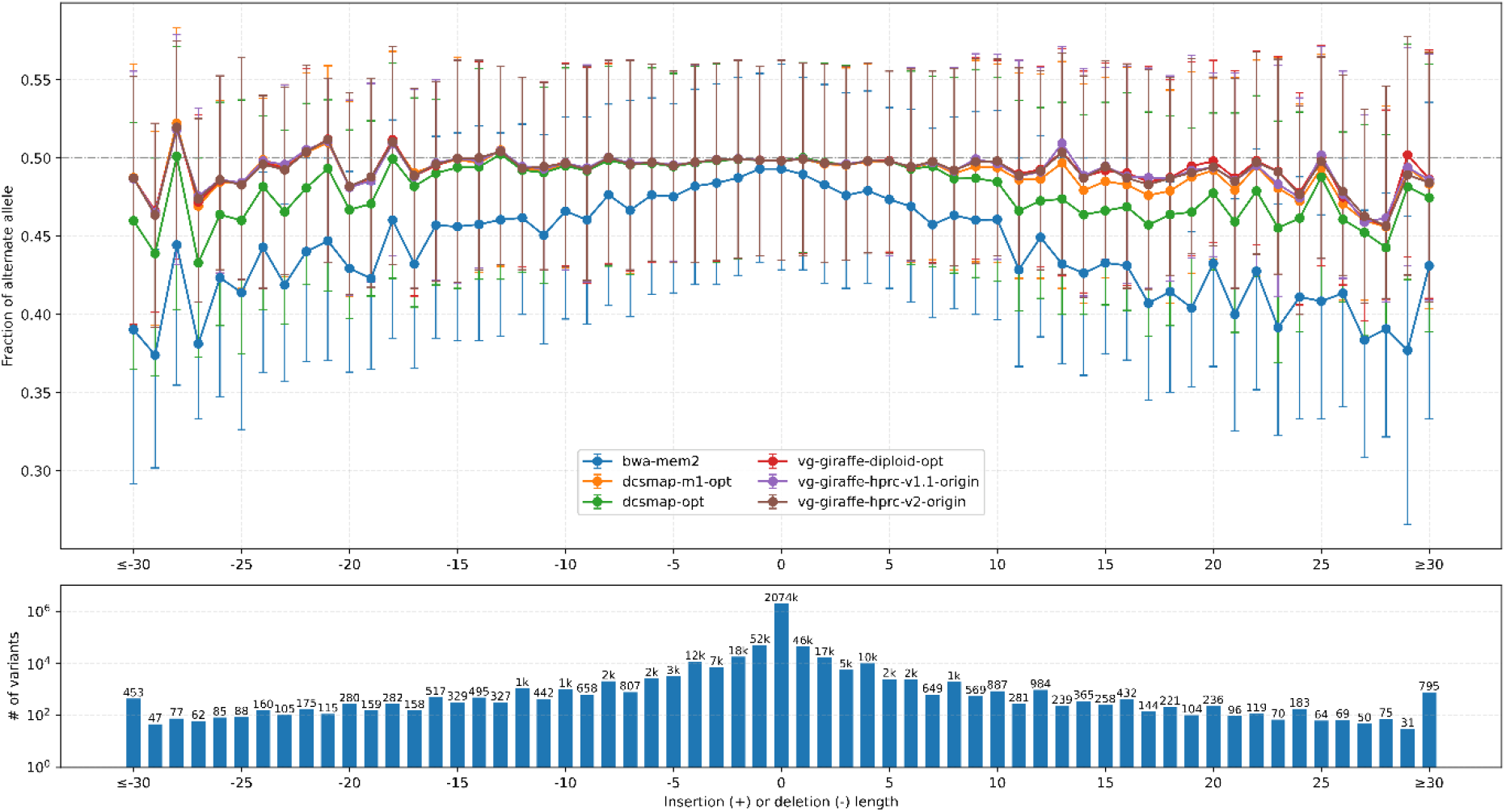
Bias-by-allele-length plots on DNBSEQ PE150 data for HG003. Variants are categorized by length: positive values for insertions, negative for deletions, and zero for SNPs at HET sites across the genome.

### Computational efficiency

Computational performance was evaluated on an Aliyun ecs.i4.8xlarge instance with 32 virtual CPU cores and 256 GiB of memory. All workflows were benchmarked using the same DNBSEQ PE150 dataset for HG003, comprising approximately 95 Gb of sequence data at ∼30× genome coverage.

The hybrid linear–graph workflow of DCSMAP substantially reduced end-to-end wall-clock time relative to pure pangenome-based workflows. dcsmap-origin completed in 3.23 h and dcsmap-m1-origin in 4.43 h, compared with 5.51 h for the personalized pangenome workflow (vg-giraffe-diploid-origin) and 5.06 and 5.51 h for the pangenome workflows using HPRC v1.1 and v2.0 filtered pangenome graph releases, corresponding to reductions of 41.4% and 19.6% for the two DCSMAP variants against the personalized pangenome workflow. CPU time followed the same ordering (76.5 and 92.4 CPU-h for the two DCSMAP variants versus 119.0–141.8 CPU-h for the pangenome-only workflows). The speed-up arises from confining graph alignment and realignment to the read subset extracted by BWA-MEM2, which reduced vg giraffe from 2.78 to 1.15 h and ABRA2 from 1.50 to 0.36 h. In addition, applying the BWA-MEM2 fast-mode alignment parameters (-S -r 3 -y 0) halved alignment time from 1.44 to 0.73 h (2.0x); these parameters skip mate rescue and reduce reseeding without modifying the aligner itself.

Because BWA-MEM2 map-and-extract runs concurrently with haplotype sampling, peak resident memory reached 129.8 GB for dcsmap-origin and 129.5 GB for dcsmap-m1-origin, both exceeding the memory available on a 32-core, 128-GB instance, whereas the graph-only workflows peaked at 69.0–88.4 GB. Resolving this bottleneck was therefore a prerequisite for deploying DCSMAP on commodity instances.

To further improve computational efficiency, we introduced software-level performance optimizations to the remaining components that preserve functionally equivalent outputs. The resulting speedups and memory changes are summarized in Figure 5. For pangenome graph haplotype sampling, vg haplotypes, vg distance index, and vg minimizer achieved 2.1 times (0.25 to 0.12 h), 3.8 times (0.39 to 0.10 h), and 2.4 times (0.39 to 0.16 h) speedups, respectively, resulting in a 2.5 times overall speedup (1.10 to 0.45 h), including an unchanged 0.07 h for k-mer counting with KMC. For graph alignment, vg giraffe improved 1.2 times (1.15 to 0.96 h). For indel realignment, ABRA2 improved 2.5 times (0.36 to 0.15 h). Because these gains came largely from improved thread efficiency rather than additional cores, CPU time fell even more steeply than wall-clock time (3.9 times for the haplotype sampling step). To enable DCSMAP to run linear alignment and haplotype sampling in parallel within a 128 GB memory, we optimized the memory usage of vg distance and vg minimizer. Their peak memory usage decreased from 88.4 to 52.1 GB (41.0% reduction) and from 68.1 to 52.5 GB (23.0% reduction), respectively, reducing the overall peak memory of haplotype sampling from 88.4 to 62.0 GB (29.8% reduction). Although ABRA2 achieved faster execution at the cost of increased memory usage (89.0 versus 31.1 GB), its memory requirement remained below 128 GB. Together these optimizations shortened dcsmap-opt to 2.17 h and dcsmap-m1-opt to 2.81 h, 32.8% and 36.6% faster than their unoptimized counterparts and 2 to 2.5 times faster than the personalized-pangenome workflow, while reducing peak memory to 104.0 and 106.7 GB. By keeping peak memory below 110 GB throughout the workflow, including the concurrent linear-alignment and haplotype-sampling stage, the optimized DCSMAP workflows could run reliably on a standard 32-core, 128-GB instance.

**Figure 5.**
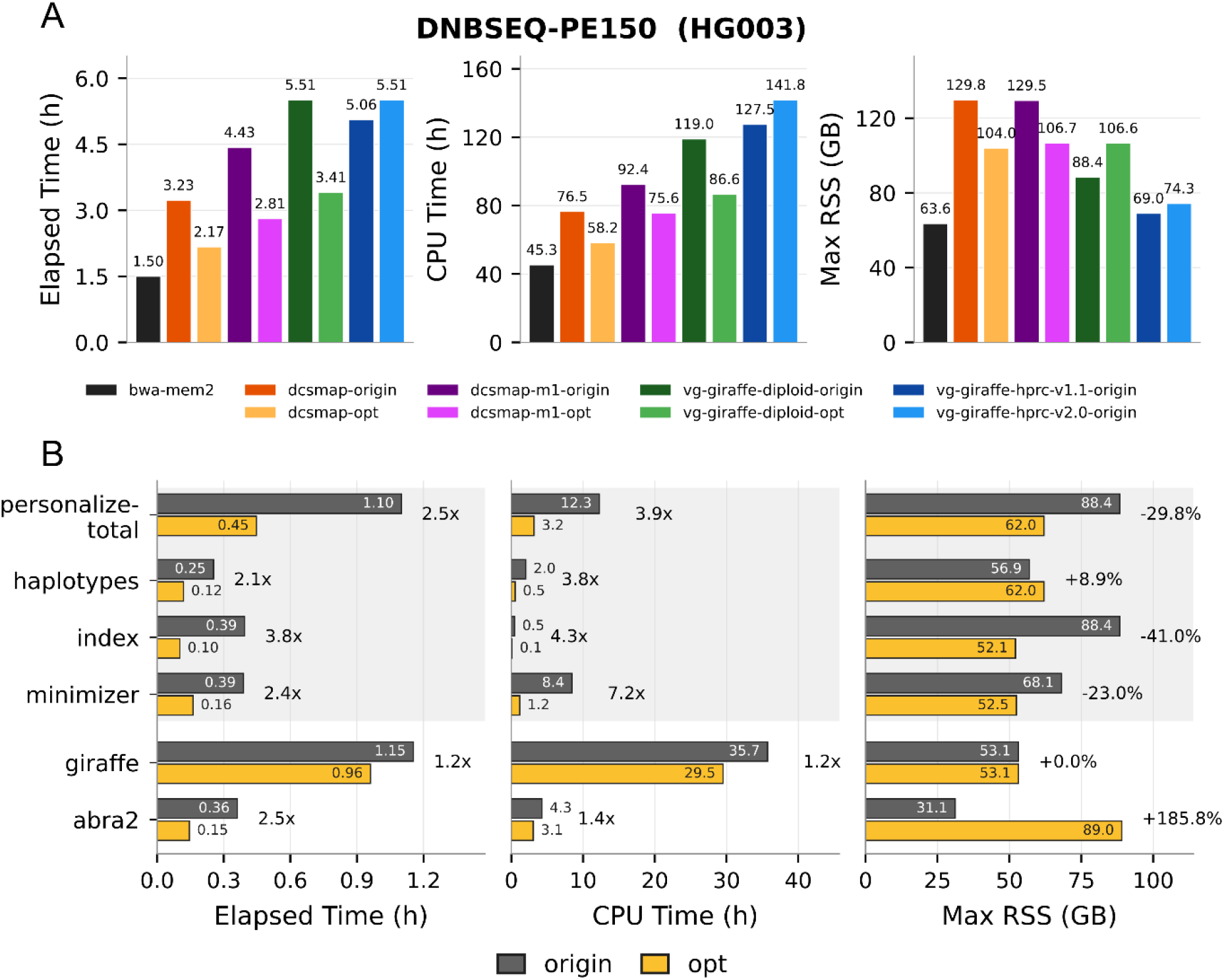
Computational performance comparison of the hybrid and other alignment workflows. Runtime and memory were measured using approximately 30× HG003 DNBSEQ PE150 data on an Aliyun ecs.i4.8xlarge instance with 32 virtual CPU cores and 256 GiB of memory. (A) End-to-end elapsed time, aggregate CPU time, and maximum resident memory size (Max RSS) for BWA-MEM2, the original and optimized dcsmap, dcsmap-m1, and personalized-pangenome workflows and the original filtered HPRC v1.1 and v2.0 population-pangenome workflows. (B) Effects of tool-level optimization on elapsed time, CPU time, and Max RSS for personalized-pangenome construction (total and its vg haplotypes, vg index, and vg minimizer components), vg giraffe alignment, and ABRA2 indel realignment. Numeric labels give the measured resource use; annotations between paired bars indicate fold speedups for elapsed and CPU time or percentage changes in Max RSS. vg giraffe and ABRA2 indel realignment were run on reads selected by dcsmap’s XGBoost classifier.

Notably, these optimizations are not specific to the hybrid architecture but apply to any haplotype-sampling-based personalized pangenome graph workflow. Applying the optimized tools to the conventional personalized-pangenome workflow reduced vg-giraffe-diploid run time from 5.51 to 3.41 h, a 38.1% reduction (1.6 times speedup), together with a decrease in CPU time from 119.0 to 86.6 CPU-h.

## Discussion

In this study, we developed DCSMAP to combine the accuracy advantages of personalized-pangenome alignment with the computational efficiency of linear alignment. DCSMAP identifies read pairs whose linear placements may be unreliable and restricts graph remapping to this subset. Across 17 evaluations covering seven GIAB samples, two sequencing platforms, and two read lengths, approximately 15% of read pairs were selected for graph remapping. Despite processing only a minority of reads with vg giraffe, the hybrid workflows consistently improved variant-calling accuracy over BWA-MEM2 and, unexpectedly, also achieved higher overall accuracy than full personalized-pangenome alignment in the primary benchmarks. These findings suggest that linear and graph alignment can play complementary roles within the same workflow.

The read-selection model is based on the observation that personalized-pangenome alignment is, on average, more accurate than alignment to a single linear reference[3,4,6]. Because validated read-level placements are unavailable for real whole-genome data, we used disagreement between BWA-MEM2 and personalized-pangenome vg giraffe alignments as a practical training label. This label is an approximation rather than read-level ground truth, as disagreement does not ensure that the graph placement is correct in every case. Nevertheless, pairing geometry and mapping quality provided broadly applicable alignment-confidence signals, and the consistent downstream improvements across samples and sequencing configurations support the effectiveness of this strategy for read routing.

The improvement over linear alignment is consistent with the ability of pangenome graphs to represent non-reference alleles and reduce reference bias. The improvement over full personalized-pangenome alignment was less expected. Although personalization reduces graph complexity, redundant or homologous paths can remain and create competing placements, lower mapping quality, or displace reads from loci supporting true variants. At false-negative sites specific to personalized-pangenome alignment in HG003, read depth and high-mapping-quality read support were lower than with DCSMAP, while most reads retained by DCSMAP originated from BWA-MEM2. These observations support a complementary division of labor: graph alignment can rescue reads affected by reference bias or alternative haplotypes, whereas linear alignment can preserve stable placements for reads already well represented by the linear reference.

The advantage of DCSMAP was smaller when HG002 was evaluated with the more comprehensive assembly-based v5.0q benchmark (Supplementary table S9) [22]. Within complex medically relevant gene regions [23], full personalized-pangenome alignment performed slightly better, mainly because DCSMAP had lower recall. Reads in repetitive, duplicated, or otherwise complex regions may receive apparently confident but incorrect linear placements and therefore escape graph remapping. Improving performance in these regions may require more diverse training data, region-aware thresholds, features describing local sequence complexity, or direct routing of reads from predefined difficult regions to the graph branch.

More generally, the central concept of DCSMAP is read routing rather than a fixed combination of linear and graph aligners. A router directs most reads to a fast alignment strategy and difficult reads to a slower but more accurate strategy. The fast branch could itself use graph alignment; for example, reads could be routed between fast and conventional vg parameter settings on the same graph. Other combinations of aligners, references, indexes, or sensitivity settings are also possible, provided that the router and its training labels are adapted to the strategies being combined.

Selective graph remapping also produced substantial computational gains. Together with optimizations to linear alignment, personalized-graph construction, graph alignment, and indel realignment, DCSMAP completed in 2.17 h compared with 5.51 h for the unoptimized full personalized-pangenome workflow and reduced peak memory sufficiently to run on a 32 core server with 128 GB of memory. The same tool-level optimizations also accelerated the conventional personalized-pangenome workflow.

In summary, DCSMAP is a practical and efficient alignment workflow that combines linear and personalized-pangenome alignment, retaining the benefits of pangenome graph alignment while substantially reducing runtime.

## Methods

### Machine-learning-based read selection

#### Training labels and data partitioning

The read-selection model was designed to identify read pairs whose linear-reference placements were potentially inaccurate or ambiguous and should therefore be remapped to a pangenome graph. Because experimentally validated read-level placements are unavailable for real whole-genome sequencing data, training labels were derived from agreement between linear-reference and personalized-pangenome alignments.

NA12891 Illumina NovaSeq PCR-free reads at approximately 30× coverage were aligned to the linear reference path of the HPRC v2.0 [19] GRCh38 [16] using BWA-MEM2 [15] with “bwa-mem2 mem -Y -S -r 3 -y 0”. The same reads were aligned using vg giraffe to a personalized pangenome graph generated by haplotype sampling from the HPRC v2.0 pangenome, followed by local indel realignment with ABRA2 [21]. The two BAM files were sorted by read name and traversed in parallel. Primary alignments for each mate were compared and assigned to one of six categories (Supplementary table S3).

Reads with concordant placements (‘SAME’) formed the negative class. Reads showing positional, indel, or missing-mate disagreement were combined into the positive class. Reads differing only in mapping quality were excluded because a difference in alignment confidence without a positional difference does not indicate which placement is correct. Unmapped reads were also excluded because the alignment-derived features required by the model, including the ‘MD’ tag, CIGAR operations, and mapping quality, were unavailable; during inference, such reads were selected unconditionally for graph remapping.

Because concordant alignments were overrepresented, the ‘SAME’ category was randomly downsampled to 12.5%, from 629,756,084 to 78,722,985 reads. After removal of the mapping-quality-only and unmapped categories, the training table contained 160,010,424 reads, comprising 49.2% negative and 50.8% positive examples. The data were divided into training, validation, and held-out test sets in an 80:10:10 ratio using stratified sampling. The test set was retained without evaluation until model configuration and hyperparameters had been finalized.

#### Alignment features and model training

Each aligned read was represented by 25 numerical features computed directly from its BWA-MEM2 BAM record, without external annotations. These features described:

- mapping quality and pairing status, including proper-pair and unmapped flags;
- insert size standardized against a library insert-size distribution estimated from the alignment stream;
- mismatch counts obtained from the MD tag, both in total and stratified into four base-quality bins;
- insertion and deletion counts and mean event lengths derived from the CIGAR string;
- soft-clipped bases;
- counts and fractions of alignment events occurring within 10 bp of a read end; and
- mean base qualities within a ±5-bp window around mismatches, insertions, deletions, and soft-clipped regions. An XGBoost classifier was trained using these features [14]. Nine hyperparameters were optimized over 80 trials by Bayesian optimization with Optuna [25]. The selected configuration used 100 trees, a maximum tree depth of 10, and a learning rate of 0.269. The final model was fitted on the training set while the validation set was used for early stopping with a patience of 30 rounds and log loss as the evaluation metric. All 100 trees were retained, during which validation log loss decreased from 0.49181 to 0.13341.

#### Streaming inference and read partitioning

Read selection was implemented in C++ as ‘extract-bam’ using the XGBoost C API. BWA-MEM2 alignment, feature extraction, classification, and BAM output were connected in a single streaming pipeline, avoiding an intermediate whole-genome alignment file. ‘extract-bam’ consumed SAM records from BWA-MEM2 through standard input, and unselected alignments were emitted as BAM records to ‘samtools sort’[26].

Alignments were grouped by read name, and each group contained both mates and all associated primary, secondary, and supplementary records. The complete read-name group was used as the decision unit. A producer thread assembled read-name groups and distributed batches of 4,000,000 records to the consumer threads. For each batch, the consumer threads parsed the records, estimated the insert-size distribution from primary paired alignments, computed the feature vectors, and scored the vectors in a batched XGBoost call. A single writer thread serialized the output.

Before feature calculation, any group containing an unmapped record was selected unconditionally and bypassed model inference. For all other groups, a read pair was selected when either mate received a classifier score greater than or equal to the default threshold of 0.50. This pair-level logical OR ensured that both mates were routed together. Selected pairs were restored to their original read orientation and written as paired, gzip-compressed FASTQ files for remapping with vg giraffe. Unselected pairs retained their BWA-MEM2 alignments and were written to a coordinate-sorted BAM file. The two outputs formed a complete, nonoverlapping partition of the input read pairs.

#### Evaluation across datasets

The deployed classifier was evaluated in 17 end-to-end runs comprising seven GIAB samples (HG001–HG007) sequenced with DNBSEQ PE150 and Illumina PE150, together with three samples (HG001, HG006, and HG007) sequenced with DNBSEQ PE100. None of these samples was used for model training. The same model, batch size, and default classification threshold were used in every run. Read-selection rates were obtained from the per-run ‘extract-bam’ counters.

### Personalized pangenome construction

A sample-specific k-mer spectrum was generated from both FASTQ files using KMC [12]. The k-mer counts, the population pangenome in GBZ format, and the corresponding haplotype index were supplied to ‘vg haplotypes’ [3]. Haplotype sampling used the observed k-mer support to select locally relevant haplotypes, retained the designated linear reference path, and generated a diploid personalized subgraph. Distance and minimizer indexes were then constructed for the sampled graph.

### Linear and graph alignment

All read pairs were first aligned to the linear reference with BWA-MEM2 [15]. The classifier retained confident linear placements and routed the selected subset to vg giraffe. Selected read pairs were aligned against the sample-specific GBZ graph using its distance and minimizer indexes. giraffe alignments were emitted as BAM records projected onto designated GRCh38 reference paths, and graph-specific path prefixes were removed to harmonize sequence names with the linear-reference BAM. Read-group information and an alignment-source tag were added to preserve sample metadata and distinguish graph-derived from linear-derived records.

For the full personalized-pangenome comparator, all read pairs rather than only the classifier-selected subset were aligned to the same type of sample-specific graph. Additional comparators included BWA-MEM2 alignment to the linear reference and vg giraffe alignment to the filtered population pangenome without sample-specific haplotype sampling. The same downstream reference coordinates and variant-calling framework were used wherever possible.

### Alignment integration and indel realignment

Alignments were locally realigned with ABRA2 [21] in candidate indel regions identified from the input BAM.

Two integration orders were evaluated. In the primary dcsmap workflow, graph-derived alignments were realigned before being merged with the retained BWA-MEM2 alignments. In dcsmap-m1, graph-derived and linear-derived alignments were first merged and the combined BAM underwent a single realignment pass. BAM files were coordinate sorted and indexed, and merging was performed while preserving compatible read-group and reference-sequence metadata.

### Small-variant calling

#### DeepVariant

Small variants were called with DeepVariant v1.10.0 using the WGS model [20]. The small model was enabled, as it is by default for the v1.10 WGS model. DeepVariant therefore used a hybrid inference strategy in which eligible, high-confidence candidates were resolved by the lightweight small model, while candidates not accepted at the small-model confidence threshold were evaluated by the full pileup-image convolutional neural network. The same pretrained WGS model was used for all alignment workflows.

The ‘make_examples’ configuration differed according to the source of the alignments. BWA-MEM2 linear-reference BAMs were analyzed with the standard WGS ‘make_examples’ settings. For dcsmap and the vg pangenome workflows, ‘make_examples’ was configured to retain mapping-quality-zero reads, use the legacy allele-counter behavior, and normalize input reads (min_mapping_quality=0, keep_legacy_allele_counter_behavior=true, and normalize_reads=true). Other DeepVariant model settings were unchanged, and only variants marked ‘PASS’ were used in downstream comparisons.

#### GATK

The linear-reference GATK workflow [27] used coordinate-sorted BWA-MEM2 alignments followed by duplicate marking. Base-quality score recalibration was performed with established known-sites resources when matching resources were available. GATK HaplotypeCaller was run in gVCF mode by chromosome, the per-chromosome gVCFs were gathered and genotyping for each sample was performed with GenotypeGVCFs.

### Variant-calling evaluation

Small-variant calls were evaluated with hap.py v0.3.12 [17]. For the primary analysis, query VCFs were compared with the GIAB v4.2.1 benchmark calls within the corresponding GIAB high-confidence regions [18,28]. HG002 was additionally assessed against the assembly-based GIAB v5.0q benchmark, including analyses restricted to complex medically relevant gene(CMRG) regions [23]. SNP and indel precision, recall, and F1 scores were obtained from the standardized hap.py comparison output. The total number of errors was defined as the sum of false-positive and false-negative calls, and relative error reduction was calculated against the specified comparator workflow. Differences between original and optimized implementations were assessed across matched workflow–dataset pairs using a two-sided paired t-test.

### Evaluating Allelic balance at HETs

Reference bias at heterozygous sites was quantified using biastools in real-read analysis mode [24]. For each alignment workflow, the coordinate-sorted BAM, the matching reference sequence, and the GIAB v4.2.1 benchmark genotypes were analyzed. Heterozygous sites were taken from the GIAB v4.2.1 truth VCF of the corresponding sample, and biastools computed allelic balance at each site, defined as the number of ALT-supporting reads divided by the total number of ALT- and REF-supporting reads mapped to that site, using its context-aware read-assignment algorithm. Only sites within the GIAB high-confidence regions were considered. To visualize the distribution of reference bias, variants were stratified by allele length: SNVs were assigned a length of zero and appear at the center of the plot, deletions extend to the left and insertions to the right with larger variants further from the center, and variants longer than 30 bp were grouped into the outermost bins. For each length bin, the mean allelic balance is reported as a central marker with the 25th and 75th percentiles shown as ticks, and the number of variants per bin is shown in a lower panel. An allelic balance near 0.5 indicates minimal reference bias, whereas lower values indicate preferential mapping of the reference allele.

The analysis was performed for HG003 on DNBSEQ PE150 (Figure 4) and Illumina PE150 (Supplementary Figure S7) data and for HG001 on DNBSEQ PE100 data (Supplementary Figure S8).

### Computational-performance evaluation

Runtime and memory benchmarks were performed on an Aliyun ecs.i4.8xlarge instance with 32 virtual CPU cores and 256 GiB of memory using approximately 30× DNBSEQ PE150 data for HG003. All workflows used the same thread count and input data. Wall-clock time, aggregate CPU time, and maximum resident memory were recorded for individual processes and for the end-to-end workflow. Original and optimized implementations of personalized-graph construction, vg giraffe alignment, and ABRA2 realignment were evaluated with otherwise matched parameters. Because linear alignment and personalized-graph construction run concurrently in dcsmap, workflow peak memory was defined as the maximum combined resident memory of simultaneously active processes rather than the largest isolated process alone.

## Availability of source code and requirements

- Project name: dcsmap
- Project home page: https://github.com/BGIResearch/dcsmap
- Operating system(s): Linux
- Programming language: C++
- Other requirements: Java 1.8.
- License: BSD 2-Clause License.
- Restrictions: dcsmap is part of the commercial software suite DCSTools and depends on DCSTools component tools to run [29]; a trial can be requested via the DCSTools official website [30].

## Data availability

All data analyzed in this study are publicly available. Download links for the HPRC v1.1 and v2.0/v2.0-eval pangenome graphs are provided in Supplementary Table S12, and those for the DNBSEQ and Illumina sequencing data from GIAB samples HG001–HG007 and NA12891 are provided in Supplementary Table S13.

## Abbreviations

ALT: alternative allele
AUPRC: area under the precision–recall curve
AUROC: area under the receiver operating characteristic curve
BAM: Binary Alignment/Map
bp: base pair
CIGAR: Concise Idiosyncratic Gapped Alignment Report
CMRG: complex medically relevant gene regions
GIAB: Genome in a Bottle
gVCF: genomic Variant Call Format
HET: heterozygous
HPRC: Human Pangenome Reference Consortium
indel: insertion or deletion
ML: machine learning
MQ: mapping quality
PCR: polymerase chain reaction
PE: paired-end
SAM: Sequence Alignment/Map
SD: standard deviation
SNP: single-nucleotide polymorphism
SNV: single-nucleotide variant
VCF: Variant Call Format
WGS: whole-genome sequencing

## Competing interests

The authors are all employees of BGI-Research.

## Funding

This research was supported by the BGI-Research and did not receive any external funding.

## Authors’ contributions

Yuxiang Li and Yong Zhang supervised the research. Qi Yang conceptualized the software. Qi Yang, and Ruiwen Wan implemented the software code. Qi Yang and Chun Gong analyzed the data. Qi Yang and Shengkang Li contributed to the writing of the manuscript. All the authors read and approved the final manuscript.

## Acknowledgements

We would like to thank DCS Cloud (https://cloud.stomics.tech/) for providing the computational resources and software support necessary for this study.

## Notes

### Competing Interest Statement

The authors have declared no competing interest.

## References

1. Degner JF, Marioni JC, Pai AA, Pickrell JK, Nkadori E, Gilad Y, et al.. Effect of read-mapping biases on detecting allele-specific expression from RNA-sequencing data. Bioinformatics. Oxford University Press (OUP); 2009; doi: 10.1093/bioinformatics/btp579.

2. Ballouz S, Dobin A, Gillis JA. Is it time to change the reference genome? Genome Biology. Springer Science and Business Media LLC; 2019; doi: 10.1186/s13059-019-1774-4.

3. Garrison E, Sirén J, Novak AM, Hickey G, Eizenga JM, Dawson ET, et al.. Variation graph toolkit improves read mapping by representing genetic variation in the reference. Nature Biotechnology. Springer Science and Business Media LLC; 2018; doi: 10.1038/nbt.4227.

4. Liao W-W, Asri M, Ebler J, Doerr D, Haukness M, Hickey G, et al.. A draft human pangenome reference. Nature. 2023; doi: 10.1038/s41586-023-05896-x.

5. Pritt J, Chen N-C, Langmead B. FORGe: prioritizing variants for graph genomes. Genome Biology. Springer Science and Business Media LLC; 2018; doi: 10.1186/s13059-018-1595-x.

6. Sirén J, Eskandar P, Ungaro MT, Hickey G, Eizenga JM, Novak AM, et al.. Personalized pangenome references. Nature Methods. Springer Science and Business Media LLC; 2024; doi: 10.1038/s41592-024-02407-2.

7. Vaddadi K, Lin M-J, Majidian S, Mun T, Langmead B. Minimizing reference bias with an imputed personalized reference. Genome Research. Cold Spring Harbor Laboratory; 2026; doi: 10.1101/gr.280989.125.

8. Grytten I, Rand KD, Nederbragt AJ, Sandve GK. Assessing graph-based read mappers against a baseline approach highlights strengths and weaknesses of current methods. BMC Genomics. Springer Science and Business Media LLC; 2020; doi: 10.1186/s12864-020-6685-y.

9. Jiang Z, Pan W, Gao R, Hu H, Gao W, Zhou M, et al.. Reference-Guided Chromosome-by-Chromosome de novo Assembly at Scale Using Low-Coverage High-Fidelity Long-Reads with HiFiCCL. Advanced Science. Wiley; 2025; doi: 10.1002/advs.202515308.

10. Sentieon Inc: Typical usage for DNAscope Pangenome. https://support.sentieon.com/docs/Pangenome_usage/pangenome/ (2025). Accessed 2026 Sept 10.

11. Li H. Minimap2: pairwise alignment for nucleotide sequences. Bioinformatics. Oxford University Press (OUP); 2018; doi: 10.1093/bioinformatics/bty191.

12. Kokot M, Długosz M, Deorowicz S. KMC 3: counting and manipulating k -mer statistics. Bioinformatics. Oxford University Press (OUP); 2017; doi: 10.1093/bioinformatics/btx304.

13. Sirén J, Monlong J, Chang X, Novak AM, Eizenga JM, Markello C, et al.. Pangenomics enables genotyping of known structural variants in 5202 diverse genomes. Science. 2021; doi: 10.1126/science.abg8871.

14. Chen T, Guestrin C. XGBoost. ACM;

15. Vasimuddin Md, Misra S, Li H, Aluru S. Efficient Architecture-Aware Acceleration of BWA-MEM for Multicore Systems. IEEE;

16. Schneider VA, Graves-Lindsay T, Howe K, Bouk N, Chen H-C, Kitts PA, et al.. Evaluation of GRCh38 and de novo haploid genome assemblies demonstrates the enduring quality of the reference assembly. Genome Research. Cold Spring Harbor Laboratory; 2017; doi: 10.1101/gr.213611.116.

17. Krusche P, Trigg L, Boutros PC, Mason CE, De La Vega FM, Moore BL, et al.. Best practices for benchmarking germline small-variant calls in human genomes. Nature Biotechnology. Springer Science and Business Media LLC; 2019; doi: 10.1038/s41587-019-0054-x.

18. Wagner J, Olson ND, Harris L, Khan Z, Farek J, Mahmoud M, et al.. Benchmarking challenging small variants with linked and long reads. Cell Genomics. Elsevier BV; 2022; doi: 10.1016/j.xgen.2022.100128.

19. Lucas JK, Hebbar P, Liao W-W, Macias-Velasco JF, Novak AM, Asri M, et al.. HPRC2: A human pangenome reference with near-complete coverage of common genetic variation. openRxiv; 2026; doi: 10.64898/2026.07.21.739710.

20. Poplin R, Chang P-C, Alexander D, Schwartz S, Colthurst T, Ku A, et al.. A universal SNP and small-indel variant caller using deep neural networks. Nature Biotechnology. 2018; doi: 10.1038/nbt.4235.

21. Mose LE, Perou CM, Parker JS. Improved indel detection in DNA and RNA via realignment with ABRA2. Bioinformatics. Oxford University Press (OUP); 2019; doi: 10.1093/bioinformatics/btz033.

22. Hansen NF, Dwarshuis N, Ji HJ, Rhie A, Loucks H, Logsdon GA, et al.. A complete diploid human genome benchmark for personalized genomics. Cell. Elsevier BV; 2026; doi: 10.1016/j.cell.2026.06.016.

23. Wagner J, Olson ND, Harris L, McDaniel J, Cheng H, Fungtammasan A, et al.. Curated variation benchmarks for challenging medically relevant autosomal genes. Nature Biotechnology. Springer Science and Business Media LLC; 2022; doi: 10.1038/s41587-021-01158-1.

24. Lin M-J, Iyer S, Chen N-C, Langmead B. Measuring, visualizing, and diagnosing reference bias with biastools. Genome Biology. Springer Science and Business Media LLC; 2024; doi: 10.1186/s13059-024-03240-8.

25. Akiba T, Sano S, Yanase T, Ohta T, Koyama M. Optuna. ACM;

26. Li H, Handsaker B, Wysoker A, Fennell T, Ruan J, Homer N, et al.. The Sequence Alignment/Map format and SAMtools. Bioinformatics. Oxford University Press (OUP); 2009; doi: 10.1093/bioinformatics/btp352.

27. McKenna A, Hanna M, Banks E, Sivachenko A, Cibulskis K, Kernytsky A, et al.. The Genome Analysis Toolkit: A MapReduce framework for analyzing next-generation DNA sequencing data. Genome Research. 2010; doi: 10.1101/gr.107524.110.

28. Zook JM, McDaniel J, Olson ND, Wagner J, Parikh H, Heaton H, et al.. An open resource for accurately benchmarking small variant and reference calls. Nature Biotechnology. Springer Science and Business Media LLC; 2019; doi: 10.1038/s41587-019-0074-6.

29. Gong C, Yuan D, Zhao Z, Li X, Chen Y, Yang Q, et al.. DCS Tools: A high-performance, resource-efficient and scalable computing suite for population-scale genomic analysis and data compression. openRxiv; 2026; doi: 10.64898/2026.03.13.711253.

30. BGI Research: DCS Tools - 高效准确的生物信息数据分析工具. https://tools.dcs.cloud/ (2026). Accessed 2026 Sept 10.

