## Supplementary Material for "DCSMAP: A Hybrid Linear-Pangenome Framework for Accurate and Efficient Read Alignment"

#### 1. Supplementary Figures

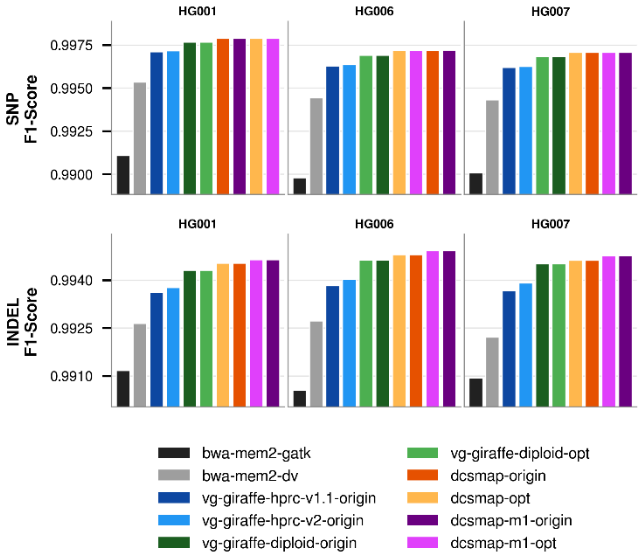


**Supplementary Figure S1.** SNP and indel F1 scores on DNBSEQ PE100 data evaluated against GIAB v4.2.1.

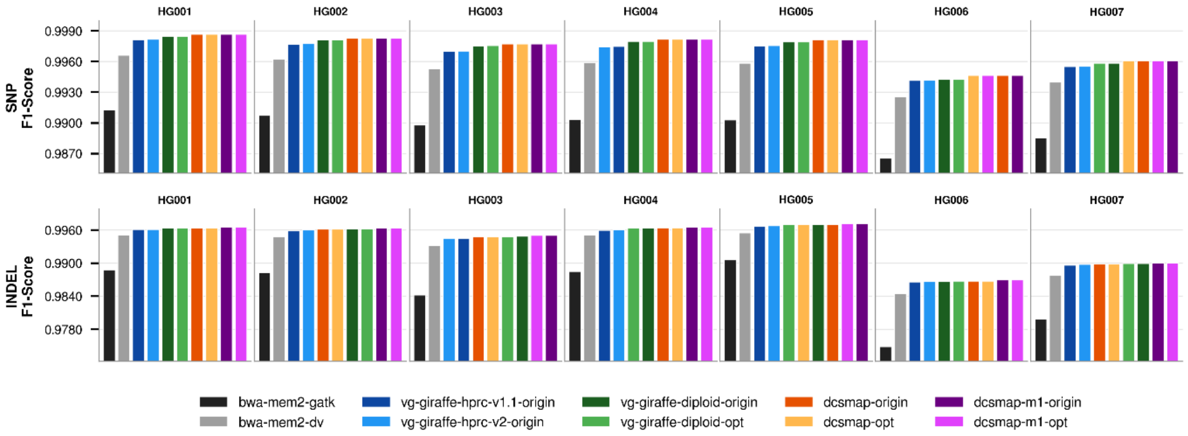


**Supplementary Figure S2.** SNP and indel F1 scores on Illumina PE150 data evaluated against GIAB v4.2.1.

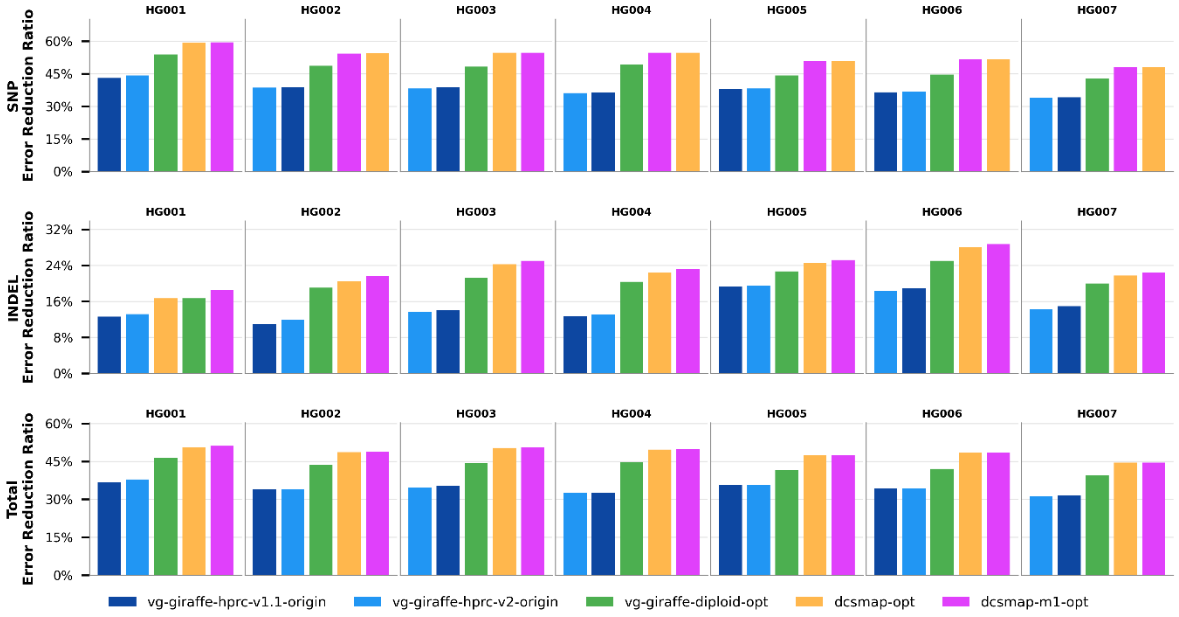


**Supplementary Figure S3.** Reduction in SNP and INDEL calling errors relative to BWA-MEM2–DeepVariant on DNBSEQ PE150 data. Total errors were calculated as the sum of false-positive and false-negative calls reported by hap.py within the GIAB v4.2.1 high-confidence regions, positive values indicate fewer errors than the BWA-MEM2–DeepVariant baseline. Error reductions are shown for each workflow and GIAB sample, with SNPs and INDELs displayed separately.

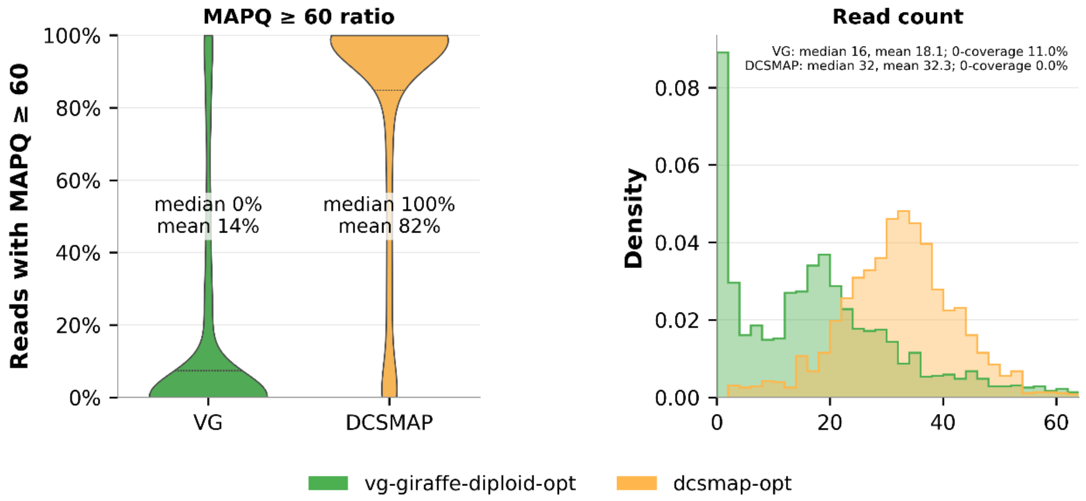


**Supplementary Figure S4.** Read support and mapping confidence at variants missed by the vg personalized pangenome workflow on DNBSEQ PE150 data for HG003. Mapping statistics were compared at 1,793 false-negative sites for vg-giraffe-diploid-opt (green) and dcsmap-opt (orange). Unmapped, secondary, duplicate, and supplementary alignments were excluded from the analysis. **Left:** Distribution of the per-site fraction of qualifying reads with mapping quality (MAPQ) $\geqslant$ 60. Sites without qualifying reads were assigned a ratio of zero. The median ratios were 0 and 1.00 for vg-giraffe-diploid-opt and dcsmap-opt, respectively (means, 0.14 and 0.82). **Right:** Distribution of qualifying read counts overlapping each variant-centered window (+-5bp).

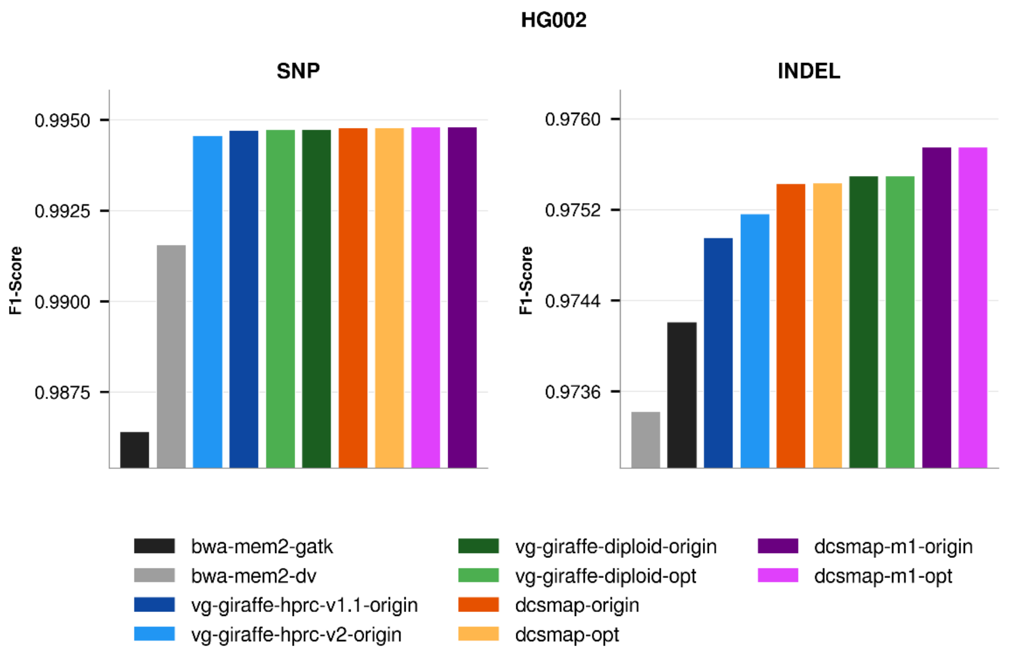


**Supplementary Figure S5.** SNP and indel F1 scores on DNBSEQ PE150 data evaluated against GIAB HG002 v5.0q.

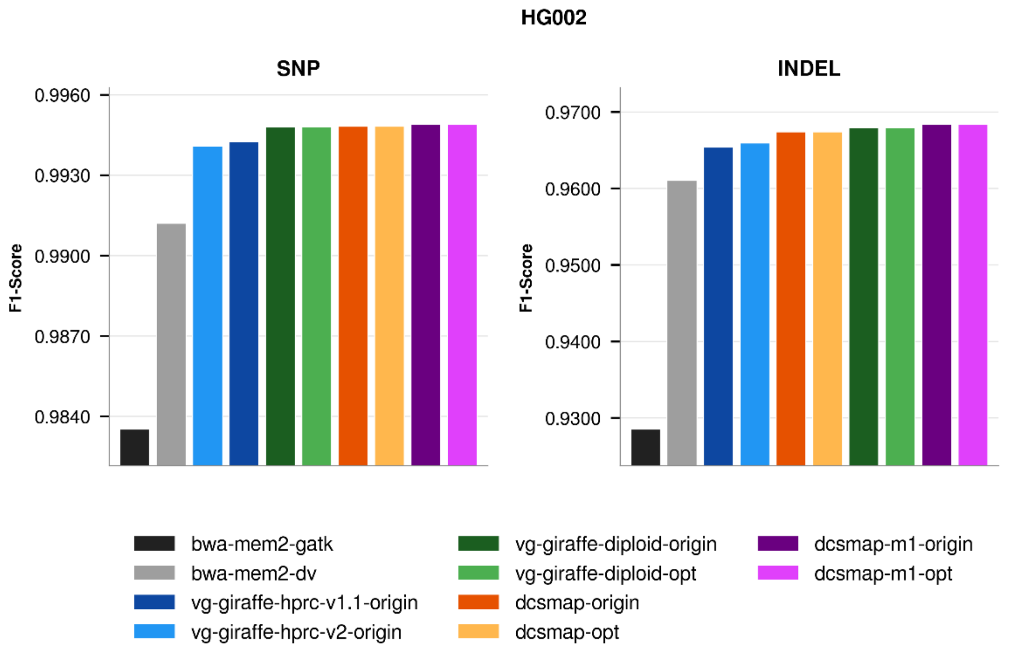


**Supplementary Figure S6.** SNP and indel F1 scores on Illumina PE150 data evaluated against GIAB HG002 v5.0q.

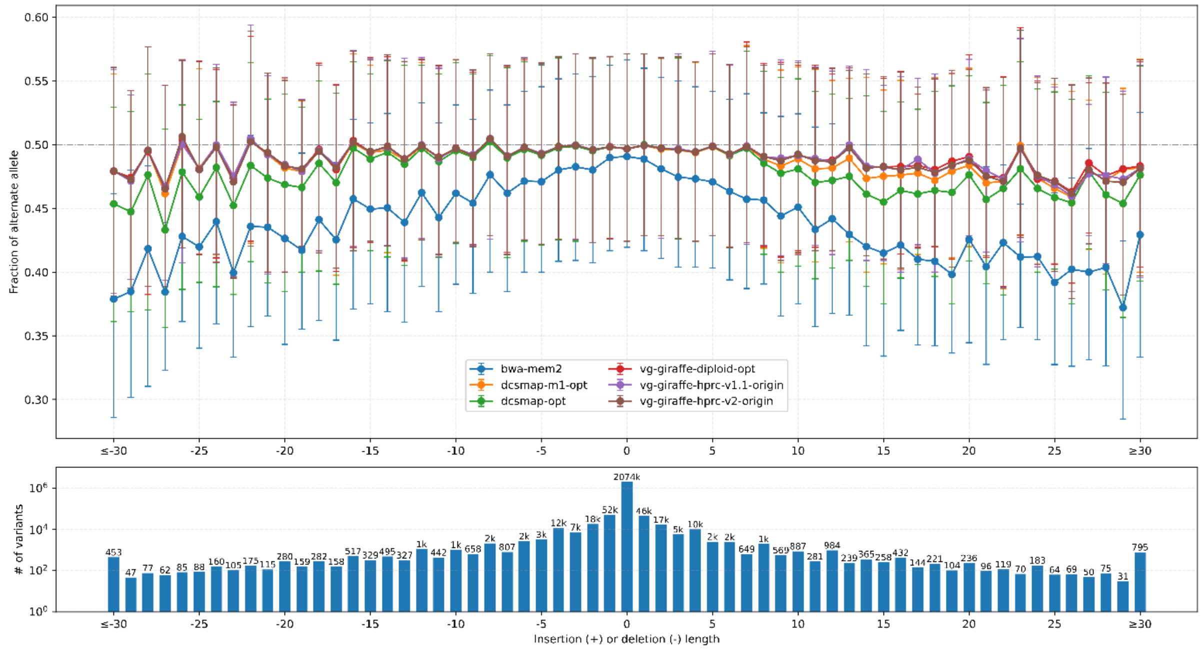


**Supplementary Figure S7.** Bias-by-allele-length plots on Illumina PE150 data for HG003. Variants are categorized by length: positive values for insertions, negative for deletions, and zero for SNPs at HET sites across the genome.

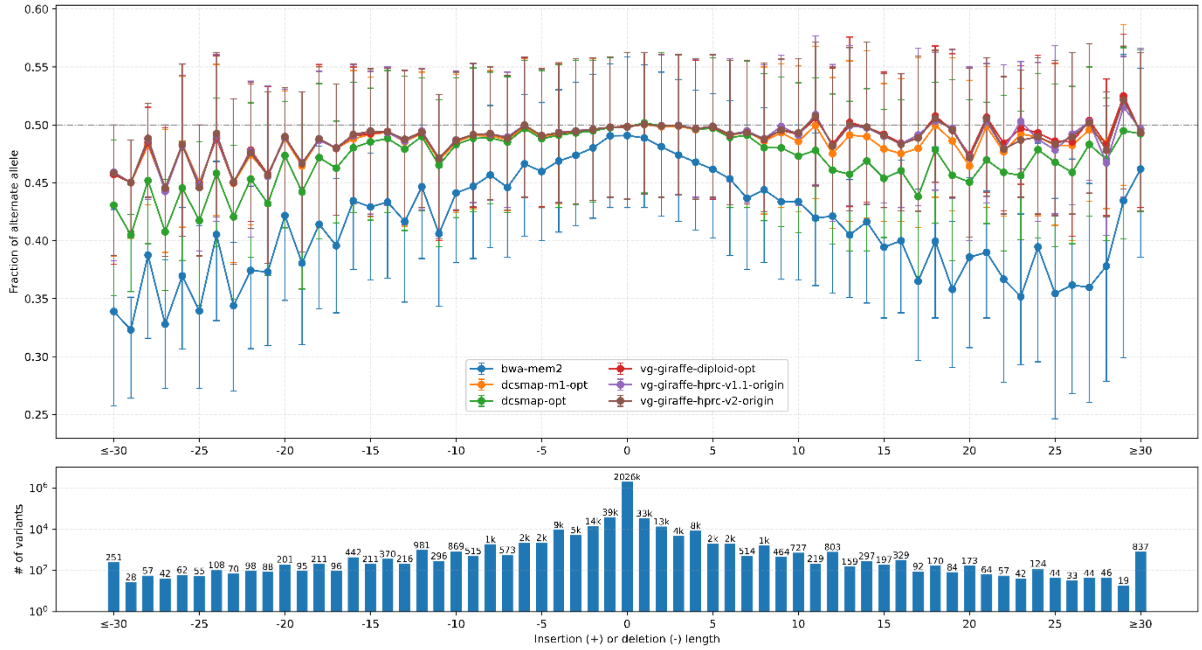


**Supplementary Figure S8.** Bias-by-allele-length plots on DNBSEQ PE100 data for HG001. Variants are categorized by length: positive values for insertions, negative for deletions, and zero for SNPs at HET sites across the genome.

#### 2. Supplementary Tables

**Supplementary Table S1. Performance of the XGBoost classifier.**

| Metric | Validation | Test |
|---|---|---|
| AUROC | 0.9831 | 0.9831 |
| AUPRC | 0.9853 | 0.9853 |
| Precision | 0.9639 | 0.9641 |
| Recall | 0.9598 | 0.9598 |
| F1 | 0.9618 | 0.9619 |
| Accuracy | 0.9613 | 0.9614 |
| False-positive rate | 0.0371 | 0.0369 |
| False-negative rate | 0.0402 | 0.0402 |
| n reads | 16,001,042 | 16,001,043 |

**Supplementary Table S2. Fraction of read pairs selected for remapping with vg giraffe.**

| Platform | n samples | Mean (%) | Median (%) | SD (%) | Range (%) |
|---|---|---|---|---|---|
| DNBSEQ-PE100 | 3 | 14.13 | 14.13 | 0.04 | 14.09–14.16 |
| DNBSEQ-PE150 | 7 | 13.23 | 12.58 | 1.55 | 12.12–16.53 |
| Illumina-PE150 | 7 | 16.65 | 15.85 | 1.91 | 15.02–19.87 |
| All runs | 17 | 14.80 | 14.16 | 2.22 | 12.12–19.87 |

**Supplementary Table S3. Concordance between the linear and personalized-pangenome alignments for NA12891.**

| Class | Criterion | Reads | % of reads | Role in training |
|---|---|---|---|---|
| SAME (0) | contig, strand, position, mapping quality and indel content all concordant | 629,756,084 | 84.13 | negative |
| POS_DIFFERENT (1) | different contig or strand, or reference offset > 100 bp | 41,959,824 | 5.61 | positive |
| MAPQ_DIFFERENT (2) | positions concordant, mapping qualities differ by > 10 | 36,084,828 | 4.82 | excluded (ambiguous) |
| INDEL_DIFFERENT (3) | graph alignment recovers a longer maximal or total indel | 4,070,809 | 0.54 | positive |
| BWA_UNMAP (4) | linear aligner placed no alignment | 1,458,393 | 0.19 | excluded (rule-based) |
| VG_MISSING (5) | graph alignment lacks the mate | 35,256,806 | 4.71 | positive |
| Total | | 748,586,744 | 100.00 | |

**Supplementary Table S4. Workflows assessed.**

| Workflow | Reference | Workflow Description |
|---|---|---|
| bwa-mem2 | GRCh38 | Maps all reads to the linear GRCh38 reference with BWA-MEM2, followed by standard BAM processing and DeepVariant/GATK variant calling. |
| dcsmap-origin | GRCh38, HPRC v2.0 personalized graph | Original-tool DCSMAP workflow: uses ML to select reads after BWA-MEM2 alignment, remaps selected reads to a personalized graph with vg giraffe, applies ABRA2 to graph-derived alignments, and merges them with retained linear alignments before DeepVariant variant calling. |
| dcsmap-opt | GRCh38, HPRC v2.0 personalized graph | Uses optimized versions of the vg tool suite and ABRA2; all other steps are identical to dcsmap-origin. |
| dcsmap-m1-origin | GRCh38, HPRC v2.0 personalized graph | Original-tool DCSMAP merge-first workflow: uses ML to select reads after BWA-MEM2 alignment, remaps selected reads to a personalized graph with vg giraffe, merges the linear and graph alignments, and then applies one ABRA2 realignment pass before DeepVariant variant calling. |
| dcsmap-m1-opt | GRCh38, HPRC v2.0 personalized graph | Uses optimized versions of the vg tool suite and ABRA2; all other steps are identical to dcsmap-m1-origin. |
| vg-giraffe-diploid-origin | HPRC v2.0 personalized graph | Uses the original vg tools to construct a personalized diploid graph by haplotype sampling, aligns all reads with vg giraffe, and performs ABRA2 realignment and DeepVariant variant calling. |
| vg-giraffe-diploid-opt | HPRC v2.0 personalized graph | Uses optimized versions of the vg tool suite and ABRA2; all other steps are identical to vg-giraffe-diploid-origin. |
| vg-giraffe-hprc-v1.1-origin | HPRC v1.1 AF-filtered pangenome graph | Aligns all reads directly to the HPRC v1.1 allele-frequency-filtered population pangenome graph with the original vg giraffe workflow, followed by ABRA2 realignment and DeepVariant variant calling. |
| vg-giraffe-hprc-v2-origin | HPRC v2.0 AF-filtered pangenome graph | Aligns all reads directly to the HPRC v2.0 allele-frequency-filtered population pangenome graph with the original vg giraffe workflow, followed by ABRA2 realignment and DeepVariant variant calling. |

### 3. Key workflow commands

##### 3.1. Personalized pangenome construction

Personalized pangenome sampling and indexing 遵循 vg 的个性化泛基因组文章指导，具体命令如下

1). K-mer counting using kmc

```
mkdir -p ${KMC_OUTPUT_DIR}/kmc_tmp
printf "%s\n%s\n" "${READS_1}" "${READS_2}" > ${KMC_INPUT_LIST}

kmc \
    -k29 \
    -t${THREADS} \
    -m64 \
    -okff \
    @${KMC_INPUT_LIST} \
    ${KMC_OUTPUT_DIR}/kmc \
    ${KMC_OUTPUT_DIR}/kmc_tmp
```

2). Personalized pangenome sampling and indexing

```
vg haplotypes \
    -v 2 \
    -t ${THREADS} \
    --num-haplotypes 32 \
    --present-discount 0.9 \
    --het-adjustment 0.05 \
    --absent-score 0.8 \
    --include-reference \
    --set-reference ${REFERENCE_NAME} \
    --diploid-sampling \
    -i ${HAPLOTYPE_INDEX} \
    -k ${KMC_OUTPUT_DIR}/kmc.kff \
    -g ${PERSONALIZED_GBZ} \
    ${POPULATION_GBZ}

vg index \
    -t ${THREADS} \
    -j ${PERSONALIZED_DISTANCE_INDEX} \
    ${PERSONALIZED_GBZ}

vg minimizer \
    -p \
    -t ${THREADS} \
    -k 29 \
    -w 11 \
    --weighted \
```

```
    --save-memory \
    -o ${PERSONALIZED_MINIMIZER_INDEX} \
    -z ${PERSONALIZED_ZIPCODES} \
    -d ${PERSONALIZED_DISTANCE_INDEX} \
    ${PERSONALIZED_GBZ}
```

${REFERENCE_NAME} was 'GRCh38' for the GRCh38-coordinate graph and 'CHM13' for the CHM13-coordinate graph.

##### 3.2. Linear alignment and ML-guided read extraction for DCSMAP

```
bwa-mem2 mem \
    -S -r 3 -y 0 -Y \
    -t ${THREADS} \
    -R "@RG\tID:${SAMPLE}\tLB:${SAMPLE}\tSM:${SAMPLE}\tPL:${PLATFORM}" \
    ${LINEAR_REFERENCE} \
    ${READS_1} \
    ${READS_2} |\
mawk '/^@/ {print;next} {print $0"\tMX:Z:BWA-MEM2"}' |\
extract-bam \
    -b 4000000 \
    -t 12 \
    -p ${EXTRACTED_READ_PREFIX} \
    -M ${XGBOOST_MODEL} |\
samtools sort \
    -l 1 \
    -OBAM \
    --threads 4 \
    -m1g \
    -o ${RETAINED_LINEAR_BAM}##idx##${RETAINED_LINEAR_BAM}.bai \
    --write-index
```

extract-bam writes selected read pairs to ${EXTRACTED_READ_PREFIX}_1.fastq.gz and ${EXTRACTED_READ_PREFIX}_2.fastq.gz for pangenome based realignment using vg giraffe. Its standard output contains the unselected linear alignments.

##### 3.3. vg Giraffe alignment and projection

```
vg giraffe \
    --progress \
    --read-group "ID:${SAMPLE}\tLB:${SAMPLE}\tSM:${SAMPLE}\tPL:${PLATFORM}" \
    --sample "${SAMPLE}" \
```

```
    -b default \
    --output-format bam \
    --left-align \
    -f ${EXTRACTED_READ_PREFIX}_1.fastq.gz \
    -f ${EXTRACTED_READ_PREFIX}_2.fastq.gz \
    -Z ${PERSONALIZED_GBZ} \
    -d ${PERSONALIZED_DISTANCE_INDEX} \
    -m ${PERSONALIZED_MINIMIZER_INDEX} \
    -z ${PERSONALIZED_ZIPCODES} \
    --ref-paths ${REFERENCE_PATH_LIST} \
    -t ${THREADS} |\
samtools addreplacerg \
    -r "ID:${SAMPLE}\tLB:${SAMPLE}\tSM:${SAMPLE}\tPL:${PLATFORM}" \
    --input-fmt BAM - |\
sed "s/${REFERENCE_PATH_PREFIX}//g" |\
mawk '/^@/ {print;next} {print $0"\tMX:Z:Giraffe"}' |\
samtools sort \
    --threads 4 \
    -m1g \
    --write-index \
    -OBAM \
    -o ${GIRAFFE_BAM}##idx##${GIRAFFE_BAM}.bai
```

${REFERENCE_PATH_PREFIX} was ‘GRCh38#0#’ or ‘CHM13#0#’. For the full personalized-pangenome comparator, ‘${READS_1}’ and ‘${READS_2}’ replaced the two extracted-read inputs; the remaining Giraffe parameters were unchanged.

##### 3.4. ABRA2 indel realignment

```
ln -sf ${LINEAR_REFERENCE} ref.fasta
ln -sf ${LINEAR_REFERENCE_INDEX} ref.fasta.fai
ln -sf ${LINEAR_REFERENCE_DICTIONARY} ref.fasta.dict
ln -sf ${LINEAR_REFERENCE_DICTIONARY} ref.dict

java -Xms24g -Xmx64g -XX:+UseG1GC \
    -jar ${GATK3_JAR} \
    -T RealignerTargetCreator \
    -drf DuplicateRead \
    --disable_bam_indexing \
    -nt ${THREADS} \
    -R ref.fasta \
    -I ${REALIGNMENT_INPUT_BAM} \
    --out ${REALIGNMENT_INTERVALS}
```

```
awk -F '[:-]' 'BEGIN { OFS = "\t" } {
    if ($3 == "") {
        print $1, $2-1, $2
    } else {
        print $1, $2-1, $3
    }
}' ${REALIGNMENT_INTERVALS} > ${REALIGNMENT_TARGETS_BED}

java -Xms24g -Xmx100g -XX:+UseG1GC \
    -jar ${ABRA2_JAR} \
    --targets ${REALIGNMENT_TARGETS_BED} \
    --in ${REALIGNMENT_INPUT_BAM} \
    --out ${REALIGNED_BAM} \
    --ref ref.fasta \
    --index \
    --threads ${THREADS} \
    --gkl
```

For dcsmap, '${REALIGNMENT_INPUT_BAM}' was the projected Giraffe BAM. For dcsmap-m1, it was the merged linear and graph-derived BAM.

##### 3.5. BAM integration

```
samtools merge \
    -f -p -c \
    -@ ${THREADS} \
    --write-index \
    -OBAM \
    -o ${MERGED_BAM}##idx##${MERGED_BAM}.bai \
    ${BAM_1} \
    ${BAM_2}
```

##### 3.6. Variant Calling using deepvariant v1.10.0

DeepVariant used the WGS model with its default small model enabled. The reported workflows did not supply '--disable_small_model=true'.

```
singularity exec \
    -B ${BIND_PATH} \
    -B ${PWD} \
    --workdir ${PWD} \
    ${DEEPVARIANT_IMAGE} \
    /opt/deepvariant/bin/run_deepvariant \
```

```
    --model_type=WGS \
    --ref ${LINEAR_REFERENCE} \
    --reads input.bam \
    --intermediate_results_dir ${DEEPVARIANT_OUTPUT_DIR}/intermediate_results \
    --output_vcf ${DEEPVARIANT_OUTPUT_DIR}/${SAMPLE}.deepvariant.vcf.gz \
    --output_gvcf ${DEEPVARIANT_OUTPUT_DIR}/${SAMPLE}.deepvariant.g.vcf.gz \
    --num_shards=${THREADS} \
    --logging_dir=${DEEPVARIANT_OUTPUT_DIR}/logs \
    --
make_examples_extra_args="min_mapping_quality=0,keep_legacy_allele_counter_behavior
=true,normalize_reads=true"
```

dcsmap, dcsmap-m1, vg-giraffe-diploid, and vg-giraffe-hprc supplied three additional ‘make_examples’ parameters: ‘min_mapping_quality=0,keep_legacy_allele_counter_behavior=true,normalize_reads=true’

The linear-reference workflow used the same command without additional ‘make_examples’ parameters.

##### 3.7. Variant calling using GATK v4.6.2

Linear alignment and duplicate marking:

```
bwa-mem2 mem \
    -t ${THREADS} \
    -R "@RG\tID:${SAMPLE}\tSM:${SAMPLE}\tPL:ILLUMINA\tLB:lib1" \
    ${LINEAR_REFERENCE} \
    ${READS_1} \
    ${READS_2} |\
samtools sort \
    -@ ${THREADS} \
    -m1g \
    -OBAM \
    -o ${SORTED_BAM}##idx##${SORTED_BAM}.bai \
    --write-index

mkdir -p ${MARKDUP_TEMP_DIR}

sambamba markdup \
    -t ${THREADS} \
    --tmpdir=${MARKDUP_TEMP_DIR} \
    ${SORTED_BAM} \
    ${MARKDUP_BAM}
```

Base-quality score recalibration:

```
${GATK} BaseRecalibrator \
    -R ${LINEAR_REFERENCE} \
    -I ${MARKDUP_BAM} \
    ${KNOWN_SITES_ARGUMENTS} \
    -O ${RECALIBRATION_TABLE}

${GATK} ApplyBQSR \
    -R ${LINEAR_REFERENCE} \
    -I ${MARKDUP_BAM} \
    --bqsr-recal-file ${RECALIBRATION_TABLE} \
    -O ${BQSR_BAM}

sambamba index \
    -t ${THREADS} \
    ${BQSR_BAM}
```

HaplotypeCaller was run separately for each reference interval:

```
${GATK} HaplotypeCaller \
    -R ${LINEAR_REFERENCE} \
    -I ${HAPLOTYPECALLER_INPUT_BAM} \
    -L ${INTERVAL} \
    -O ${SAMPLE}.${INTERVAL}.g.vcf.gz \
    -ERC GVCF \
    --native-pair-hmm-threads ${HAPLOTYPECALLER_THREADS}
```

The per-interval gVCFs were gathered in reference-dictionary order and genotyped:

```
${GATK} GatherVcfs \
    ${GVCF_INPUT_ARGUMENTS} \
    -O ${SAMPLE}.g.vcf.gz

${GATK} IndexFeatureFile \
    -I ${SAMPLE}.g.vcf.gz

${GATK} GenotypeGVCFs \
    -R ${LINEAR_REFERENCE} \
    -V ${SAMPLE}.g.vcf.gz \
    -O ${SAMPLE}.vcf.gz

${GATK} IndexFeatureFile \
    -I ${SAMPLE}.vcf.gz
```

##### 3.8. hap.py evaluation

The hap.py v0.3.12 command was:

```
hap.py \
    ${TRUTH_VCF} \
    ${QUERY_VCF} \
    -f ${HIGH_CONFIDENCE_BED} \
    -r ${LINEAR_REFERENCE} \
    -o ${HAPPY_OUTPUT_PREFIX} \
    --engine=vcfeval \
    --threads ${THREADS}
```

GIAB v4.2.1 truth calls and high-confidence regions were used for the primary GRCh38 analysis. HG002 v5.0q analyses used the matching v5.0q truth set and confident regions. CMRG analyses used the intersection of the benchmark high-confidence regions and CMRG intervals.
